# Expression of Arabidopsis Extracellular Vesicle Protein Markers in *Nicotiana benthamiana* Reveals Distinct Vesicle Subpopulations

**DOI:** 10.1101/2025.05.06.652549

**Authors:** Suchismita Ghosh, Roger W. Innes

**Affiliations:** Department of Biology, Indiana University, Bloomington, Indiana, USA

**Keywords:** extracellular vesicles, TIRF Microscopy, *Arabidopsis thaliana*, *Nicotiana benthamiana*, plant-microbe interactions

## Abstract

Mammalian extracellular vesicles (EVs) are heterogeneous in nature based on their protein content, RNA content, density, size, and functions. In contrast, our understanding of plant EV diversity is quite limited. Multiple plant EV protein markers have been identified. Two of these, TETRASPANIN 8 (TET8) and PENETRATION 1 (PEN1), appear to mark distinct subpopulations of plant EVs. To further assess the diversity of plant EV subpopulations, we purified EVs from *N. benthamiana* transiently expressing multiple EV marker proteins and then assessed colocalization of these markers using high resolution Total Internal Reflection Fluorescence Microscopy (TIRF-M). We confirmed that TET8 and PEN1 indeed mark distinct EV populations, as they colocalized only 4.7% of the time. This value was nearly identical to that found for EVs purified from transgenic Arabidopsis co-expressing these two markers, demonstrating that transient expression of Arabidopsis EV proteins in *N. benthamiana* can be used to assess EV subpopulations, bypassing the requirement of generating transgenic plants for every marker combination of interest. We then used the *N. benthamiana* system to assess colocalization of PEN1 and TET8 with the EV markers PATELLIN1 (PATL1), ANNEXIN2 (ANN2), and RPM1-INTERACTING PROTEIN4 (RIN4). PATL1 and ANN2 colocalized with PEN1 56.6% and 46.6% of the time, respectively, whereas they colocalized with TET8 only 28.4% and 30.8% of the time, respectively. In contrast to PATL1 and ANN2, the RIN4 protein colocalized with TET8 more frequently than with PEN1 (30% versus 13%). Together, these results indicate that plant EVs are heterogeneous in their protein cargos and that TET8 marks a distinct subpopulation of EVs. PEN1, PATL1, and ANN2 commonly mark the same EV population that is distinct from TET8-labeled EVs, while RIN4 more often associates with TET8-labeled EVs. These findings suggest that plants possess at least two different pathways for EV biogenesis and secretion.

## Introduction

Extracellular vesicles (EVs) are cell-derived membrane bound structures that contain cargo molecules such as proteins, lipids, and nucleic acids (Gyorgy et al., 2011; Akers et al., 2013; Rutter and Innes, 2017; Cai et al., 2018). In eukaryotic cells, there are at least three EV biogenesis pathways. EVs can be derived from direct budding of vesicles from the plasma membrane. Such vesicles are referred to as microvesicles. EVs can also be formed when a multivesicular body (MVB) containing several intraluminal vesicles (ILVs) fuses with the plasma membrane, releasing its internal components, including ILVs, into the extracellular space. The third method of EV biogenesis occurs during programmed cell death (PCD), which leads to the formation of apoptotic bodies that are formed by blebbing off the plasma membrane during PCD (Gyorgy et al., 2011; Akers et al., 2013). Cells from all three major domains of life (Archaea, Bacteria and Eukarya), including plants and animals, secrete EVs (Rutter and Innes, 2017; Gill et al., 2019).

Extracellular vesicles from mammals are highly heterogeneous in their functions, contents and biogenesis (Jeppesen et al., 2019; Mathieu et al., 2019). EVs have been implicated in adaptive immunity, by presenting antigens on their surface and facilitating immune responses (Thery et al., 2002; Giri and Schorey, 2008). EVs can also carry markers for certain cancers, and EVs derived from certain tumors can target specific organs to promote tumor metastasis (Hoshino et al., 2015). EVs serve as markers for diagnosis of diseases such as Parkinson’s and bladder cancer (Welton et al., 2010; Shi et al., 2023). EVs also carry RNAs, including microRNAs (miRNAs) and messenger RNAs (mRNAs), which can be diagnostic for malignant or benign tumors (Valadi et al., 2007; Skog et al., 2008; Pigati et al., 2010).

Plant EVs increase in number when *Arabidopsis* is infected with a bacterial pathogen (*Pseudomonas syringae* strain DC3000) or the fungal pathogens *Botrytis cinerea*, *Colletotrichum higginsianum* (Rutter and Innes, 2017; He et al., 2021; Koch et al., 2024), suggesting that they play a role in immune responses. Plant EVs have also been purified from *N. benthamiana* and *Medicago sativa* (He et al., 2021; Ghosh et al., 2024) and production of *M. sativa* EVs is also upregulated in response to a fungal infection (Ghosh et al., 2024). Consistent with a role in immunity, plant EVs appear to localize to sites of attempted fungal penetration (An et al., 2006; An et al., 2006; Micali et al., 2011).

Plant EVs may be derived from MVBs, which are often seen at infection sites, especially in the vicinity of papillae, which are extracellular depositions of polysaccharides and lipids located beneath attempted penetration sites of filamentous pathogens (Xu and Mendgen, 1994; An et al., 2006; An et al., 2006). When MVBs fuse with the plant plasma membrane, the fused structures are referred to as paramural bodies (PMBs). PMBs accumulate at the periphery of *M. sativa* epidermal cells upon attempted infection by the incompatible fungal pathogen *C. higginsianum* (Ghosh et al., 2024). EVs are upregulated during such incompatible interactions, suggesting that paramural bodies and MVBs are a source of EV biogenesis, and that there may be a class of EVs specifically upregulated during immune responses (Ghosh et al., 2024).

Density gradient studies have been successful in showing that animal EVs come in different densities and sizes, each with distinct cargo proteins and RNAs (Kowal et al., 2016; Jeppesen et al., 2019; Temoche-Diaz et al., 2019). Recently, a similar density gradient separation study was performed on *Arabidopsis* EVs and it was shown that plant EVs can be separated into high, medium and low density EVs based on their buoyant densities (Koch et al., 2024). It was also shown that the plant EV marker proteins PENETRATION 1 (PEN1) and PATELLIN 1 (PATL1) co-fractionate with all three of these EV populations, while TETRASPANIN 8 (TET8) fractionated only with the medium density EVs (Koch et al., 2024). Studies performed with transgenic *Arabidopsis* have identified at least two populations of plant EVs, one carrying TET8, and a second population carrying PEN1 (He et al., 2021; Koch et al., 2024). However, unlike the animal field, the plant field lacks a quick and robust way to determine EV subpopulations and their cargo contents. Identifying EV marker proteins that occupy different EV subpopulations will provide significant insights into the biogenesis of plant EVs and their likely roles in plant biology.

Our current lack of understanding of plant EV diversity stems, in part, from a lack of antibodies that detect native EV proteins and a lack of transgenic plants that express EV markers tagged with fluorescent proteins in pairwise combinations. Generation of the latter represents a tedious and time-consuming process. We therefore evaluated whether EV marker sorting could be studied using transient expression of EV markers in *N. benthamiana,* as transient expression is much faster and more facile than generation of transgenic plants. We also wished to study individual EVs to ensure that our conclusions were accurate and not due to technical artifacts. To reduce artifacts, we used Total Internal Reflection Fluorescence Microscopy (TIRF-M), which minimizes background signals and only images a limited specimen region immediately adjacent to the coverslip (Han et al., 2021). To minimize EV clumping, we used EGTA as a chelating agent when isolating vesicles and resuspending vesicle pellets, as it chelates ions such as Ca^2+^ that might lead to vesicle clumping (Ebrahimi and Keshtgar, 2020; Borniego et al., 2025). To further minimize artifacts caused by EV aggregation, we employed the Fiji Tool DiAna (Gilles et al., 2017; Koch et al., 2024) to determine the distance between two fluorescent EV marker signals before classifying them as occupying the same or different vesicle. Together, these measures enabled us to accurately quantify the heterogeneity of plant EVs.

We focused our analyses on PEN1, PATL1, TET8, ANNEXIN2 (ANN2), and RIN4, all of which have been shown to be markers of EVs in plants (Rutter and Innes, 2017; He et al., 2021). PEN1, also known as SYNTAXIN OF PLANTS121 (SYP121) is a SNARE protein that is involved in vesicle trafficking, membrane fusion, papillae formation and immunity in plants (Assaad et al., 2004; Nielsen et al., 2012; Waghmare et al., 2018). GFP-PEN1 protein in papillae retains its fluorescence signal despite the relatively low pH of the apoplast, which suggested that PEN1 may be packaged inside EVs (Meyer et al., 2009). Consistent with this observation, protease protection assays have shown that PEN1 is protected inside the lumen of EVs (Rutter and Innes, 2017; He et al., 2021).

PATL1 is a cell-plate associated protein, and shares sequence similarities with proteins involved in membrane trafficking (Peterman et al., 2004). PATL1 binds to phosphoinositides and also binds to PLASMODESMATA-LOCATED PROTEIN1 (PDLP1) during powdery mildew infection (Peterman et al., 2004; Caillaud et al., 2014). PATL1 is also found in the EV proteome and is protected inside the lumen of EVs (Rutter and Innes, 2017).

TET8 is a homolog of the human EV marker CD63. It has recently been shown to bind to glycosyl inositol phosphoceramides (GIPCs), which are enriched in plant EVs (Liu et al., 2024). Notably, knockout of *TET8* reduces total EVs in the apoplast, indicating that TET8 may contribute directly to biogenesis of at least a subset of plant EVs.

ANN2 is an RNA binding protein that is associated with extracellular vesicles (Rutter et al., 2017). It binds to small RNAs non-specifically (He et al., 2021). ANN2 has been proposed to localize with TET8-containing EVs, as TET8 EVs are thought to carry RNA (He et al., 2021).

RIN4 is a plant-specific protein with a large intrinsically disordered region that promotes formation of condensates (Zhu et al., 2025). It also contains two nitrate-inducible (NOI) domains that mediate interaction with the exocyst complex subunit EXO70 and a C-terminal acylation motif that targets RIN4 to the plasma membrane (Kim et al., 2005; Sabol et al., 2017; Redditt et al., 2019). These features suggest that RIN4 is involved in exocytosis. Consistent with this, overexpression of RIN4 reduces secretion of callose induced by the bacterial flagellin-derived peptide flg22 (Kim et al., 2005).

Our analyses confirmed that PEN1 and TET8 are mostly sorted into different vesicles, with very little overlap between them. We also found that PATL1 and ANN2 often mark the same vesicles as PEN1 as they colocalized with PEN1 vesicles. PATL1 and ANN2 only colocalized with TET8 EVs between 25 to 30% of the time, indicating that TET8 vesicles might have a different biogenesis process than PEN1-labelled vesicles. In support of this conclusion, the frequency of vesicles containing TET8 alone was always higher than the frequency of TET8 vesicles with either PATL1 or ANN2. Notably, RIN4 co-localized with TET8 more frequently than with PEN1, suggesting that RIN4 may function with TET8 to produce the TET8 class of vesicles. Since all EV marker protein pairs colocalized at least some of time, it suggests that the different subclasses of EVs may share steps in the EV biogenesis pathway, and/or are derived from a common source, such as the plasma membrane.

## Results

### Recombinant EV marker proteins express in *N. benthamiana* leaves

To assess colocalization of the EV marker proteins PEN1, PATL1, TET8, ANN2, and RIN4, we first generated fluorescent protein fusions with either eGFP, tGFP or mCHERRY to enable transient expression in *N. benthamiana* in all pairwise combinations. To confirm that the recombinant proteins expressed well in *N. benthamiana*, we transiently expressed them using *Agrobacterium tumefaciens* and observed their expression under fluorescence microscopy. Supplementary Figure S1A shows that the empty vector controls and buffer (MgCl_2_) control did not fluoresce in the GFP or RFP channels. Supplementary Figure S1B shows that all fusion proteins accumulated well in the epidermal cells. PATL1, PEN1 and RIN4 appeared to be mostly plasma membrane localized based on their very sharp demarcation of the cell periphery, whereas ANN2 a appeared to be mostly cytoplasmic based on broader and uneven accumulation around the cell periphery and the presence of cytoplasmic strands. TET8 was intermediate between these two patterns, appearing mostly peripheral, but with its distribution somewhat patchy.

**Supplementary Figure S1.**
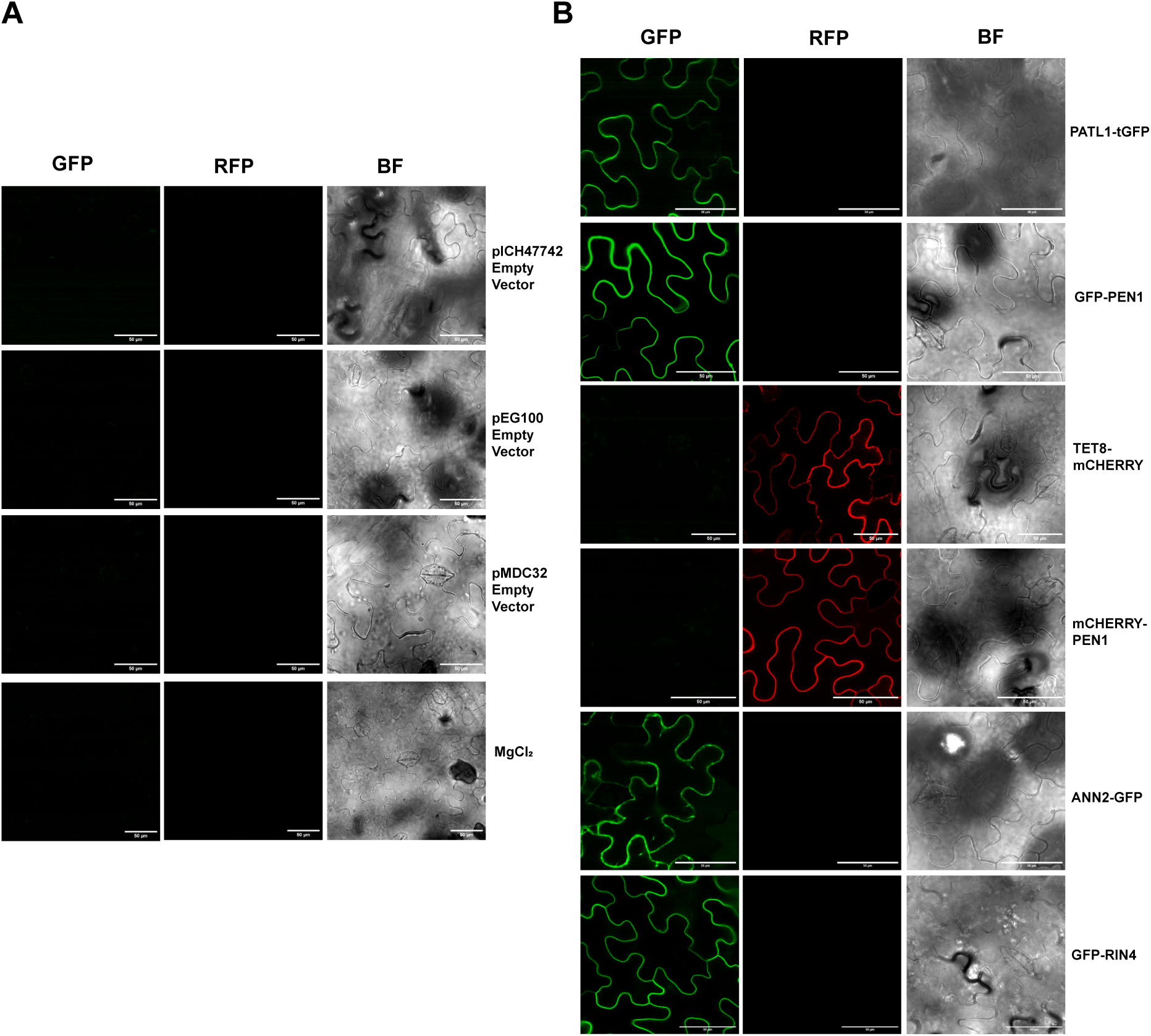
EV marker proteins are expressed in *N. benthamiana* leaves. **A,** Lack of fluorescent signal in *N. benthamiana* leaves expressing an empty vector control or injected with MgCl2 alone. **B,** Localization of recombinant EV markers expressed in pavement cells of *N. benthamiana* leaves. PEN1, TET8, ANN2 were under the control of Cauliflower Mosaic Virus 35s promoter, and PATL1 was under the control of a promoter region from Cassava Vein Mosaic Virus. All images were taken 60 hours after injecting *N. benthamiana* leaves with their respective Agrobacterium strain. Scale bar represents 50 µm.

### Recombinant EV marker proteins are fully protected inside *N. benthamiana* vesicles

We also confirmed that full-length recombinant protein for each construct is expressed using immunoblots (Supplementary Figure S2A), which indicates that the signal observed using confocal microscopy mostly corresponds to intact protein and not degradation products. We used *N. benthamiana* leaves injected with MgCl_2_ or *A. tumefaciens* carrying an empty vector as negative controls.

**Supplementary Figure S2.**
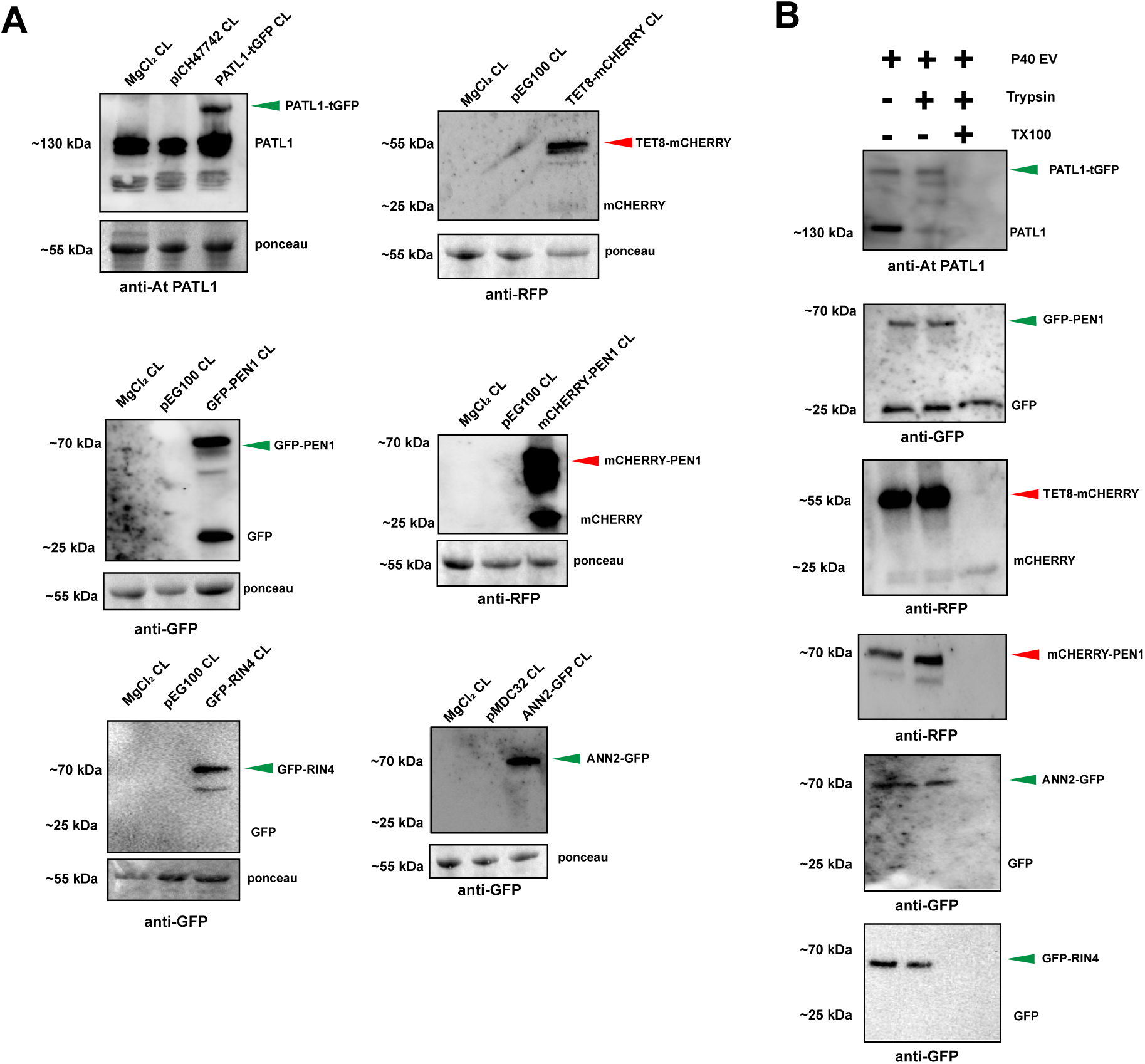
Transiently expressed EV marker proteins are packaged inside EVs in *N. benthamiana* leaves. **A,** Immunoblots validating transient expression of full-length recombinant EV marker proteins in whole cell lysate (CL). The middle lane in each blot is an empty vector control. **B,** Immunoblots of P40 EVs isolated from the apoplast of *N. benthamiana* leaves transiently expressing the indicated proteins. Full-length recombinant EV proteins are fully protected from trypsin treatment and are susceptible to trypsin digestion only when pretreated with detergent.

To confirm that these EV markers with their tags were packaged properly inside EVs, we performed a protease protection assay as described in (Rutter and Innes, 2017). In brief, we isolated P40 pellets (the pellet obtained after centrifuging apoplastic wash fluid at 40,000*g* for one hour) from *N. benthamiana* leaves expressing the recombinant proteins. We then resuspended these pellets in a small volume of buffer and treated them with protease (trypsin) for one hour. If the recombinant proteins are protected inside the lumen of the extracellular vesicles, they should be resistant to protease treatment. All recombinant EV marker proteins were resistant to trypsin digestion in the absence of detergent but were fully digested when EVs were treated with detergent prior to digestion (Supplementary Figure S2B). These results confirm that the recombinant proteins are being packaged into EVs.

### GFP-PEN1 takes 6 hours longer to accumulate in EVs than in the plasma membrane

It was not known previously how long it takes for EV associated proteins to accumulate in EVs. To address this question, we used a dexamethasone-inducible promoter system to transiently express EV markers in *N. benthamiana* and then assessed marker protein levels in EVs over a time course following induction. We first checked the expression of the GFP-PEN1 protein at three time points post dexamethasone spraying, 6h, 12h and 24h. We found that the fluorescent signal from GFP-PEN1, as observed by confocal microscopy, was strongest at 6h post dexamethasone spraying, and decreased with time, with 24h showing the weakest signal (Figure 1A). We then isolated P40 EVs from these plants and performed immunoblots to assess GFP-PEN1 protein levels. We also isolated proteins from the leaves, but the proteins were isolated after collection of the apoplastic wash to remove apoplastic GFP-PEN1. TEM images in Figure 1B show that EVs were present in P40 fractions from both GFP-PEN1 leaves as well as the empty vector control leaves. TIRF-M imaging of these P40 EVs showed fluorescent puncta only in P40 fractions from GFP-PEN1 expressing leaves and not in P40 fractions from empty vector controls (Figure 1C). The most numerous puncta were observed at the 24h timepoint. Consistent with this observation, GFP-PEN1 protein accumulation in the P40 fraction was not apparent until 12h post induction and appeared stable at 24h (Figure 1D). This result indicates that there is a delay between accumulation of GFP-PEN1 in the plasma membrane and accumulation in the apoplast, likely reflecting the time required to transit the EV secretion pathway.

**Figure 1.**
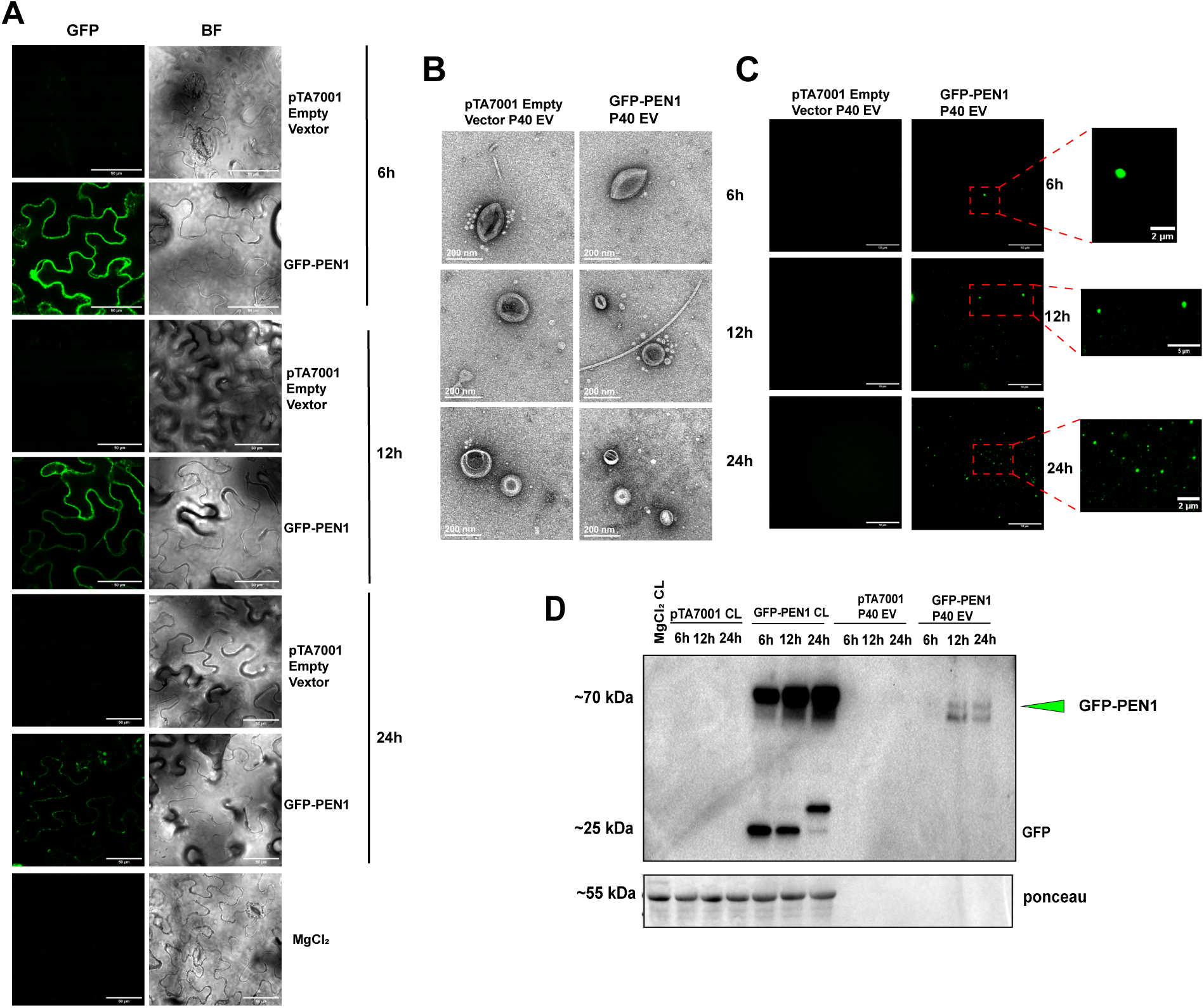
EV marker proteins accumulate in the apoplast roughly 12h after induction of expression. **A,** Confocal microscopy images of GFP-PEN1 transiently expressed in pavement cells of *N. benthamiana* leaves 6h, 12h and 24h after dexamethasone application **B,** Negative stain TEM images of P40 EVs isolated from the apoplast of *N. benthamiana* leaves 6h, 12h and 24h after dexamethasone application. **C,** TIRF-M images of P40 EVs isolated from the apoplast of *N. benthamiana* leaves 6h, 12h and 24h after dexamethasone application. Green puncta are presumed to be EVs containing GFP-PEN1. The red boxed region is zoomed in for better visualization of EVs. **D,** Immunoblot of P40 EVs isolated from *N. benthamiana* leaves transiently expressing GFP-PEN1. Ponceau staining shows EV pellets have little to no cellular contamination. Scale bar in A represents 50µm. Scale bar in B represents 200 nm. Scale bar in C represents 10 µm. Scale bar in the zoomed image of 6h and 24h in C represents 2 µm. Scale bar in the zoomed image of 12h C represents 5 µm.

### Only full-length protein gets packaged inside EVs

Before assessing colocalization of recombinant proteins in EVs using TIRF microscopy, we wished to confirm that fluorescent puncta observed by TIRF-M were coming from full-length marker proteins inside extracellular vesicles and not autofluorescence or free fluorescent proteins. Supplementary Figure S3 shows that even though free GFP and free mCherry accumulate to high levels when transiently expressed in *N. benthamiana* leaves, we could not detect a fluorescent signal in P40 pellets using TIRF-M imaging. TEM images of these P40 fractions confirmed that they contained EVs (Supplementary Figure S3B). The absence of free GFP and free mCherry in EVs was confirmed by immunoblot (Supplementary Figure S3C). As an additional negative control, we also isolated P40 pellets from *N. benthamiana* leaves injected with MgCl_2_ buffer. Supplementary Figure S3B shows that these samples did not fluoresce with TIRF-M, while TEM imaging confirmed EV isolation. These results confirmed that fluorescent puncta observed using our TIRF-M settings should correspond to EV marker proteins and not autofluorescence or free fluorescent proteins resulting from degradation.

**Supplementary Figure S3.**
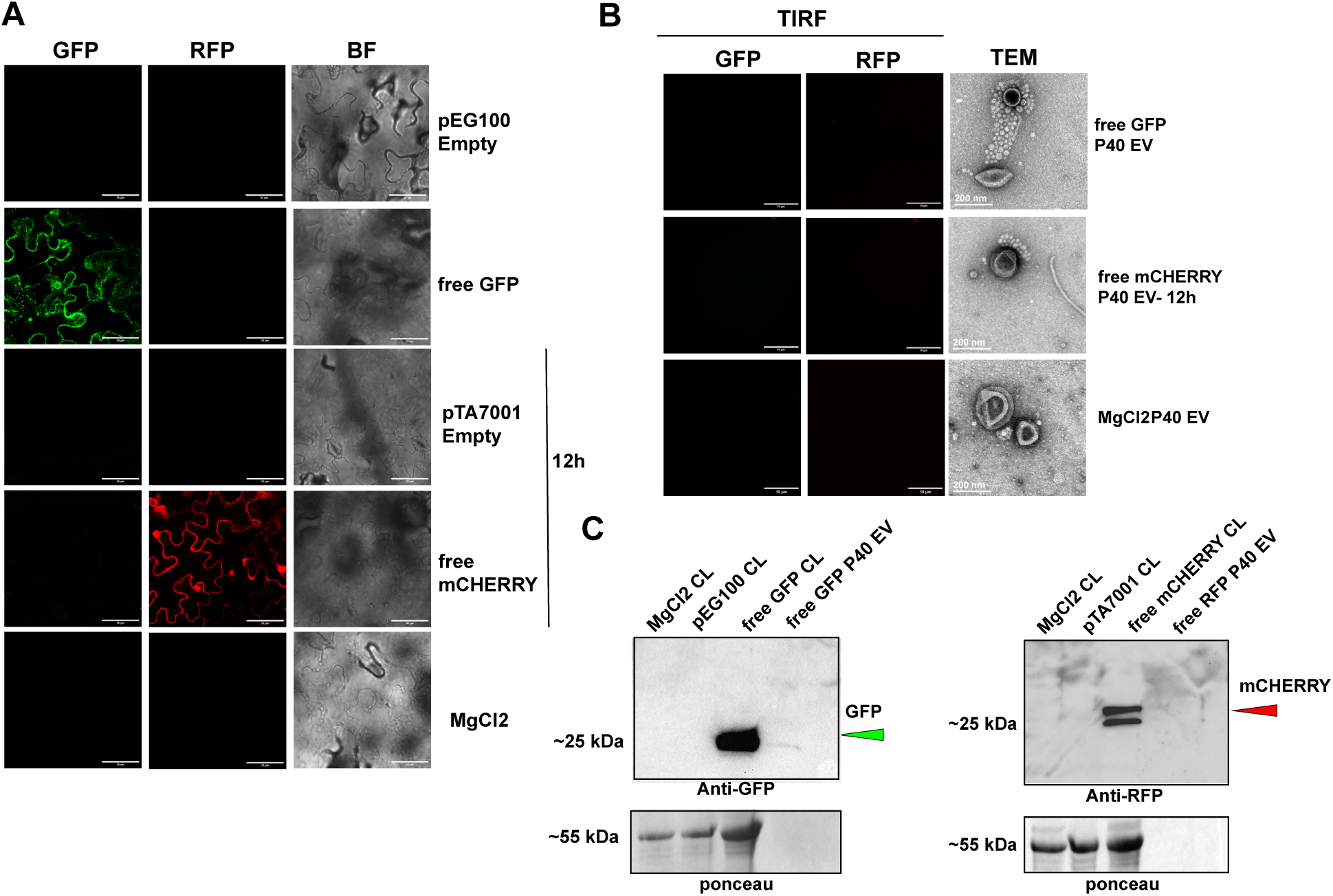
Free GFP and free mCherry do not accumulate in P40 EVs. **A,** Localization of free GFP and free mCherry transiently expressed in pavement cells of *N. benthamiana* leaves. Free GFP was expressed using pEG100 with a CaMV 35S promoter. Free mCherry was expressed using pTA7001 with a dexamethasone-inducible promoter. P40 fractions were collected 48 hrs after inoculation (free GFP) or 12 hrs after dexamethasone application (free mCherry). **B,** TIRF-M and TEM images of P40 EVs isolated from *N. benthamiana* expressing free GFP and free mCHERRY and control MgCl2 buffer. **C,** Immunoblots of cell lysates (CL) and P40 fractions isolated from *N. benthamiana* leaves expressing free GFP and free mCherry. Scale bar in panel A represents 50 µm. Scale bar in panel B TIRF images represents 10 µm. Scale bar in panel B TEM images represents 200 nm.

### PEN1 and TET8 mark distinct EV populations

Previous reports have shown that TET8 and PEN1 label distinct EV populations in Arabidopsis (He et al., 2021; Koch et al., 2024). To assess whether the *N. benthamiana* transient expression system recapitulates these findings, we co-expressed GFP-PEN1 and TET8-mCherry in *N. benthamiana* leaves. As seen in Supplementary Figure S4A, most cells expressed both genes. We confirmed the isolation of EVs using negative stain TEM, which revealed cup-shaped structures typical of EVs (Supplementary Figure S4B).

**Supplementary Figure S4.**
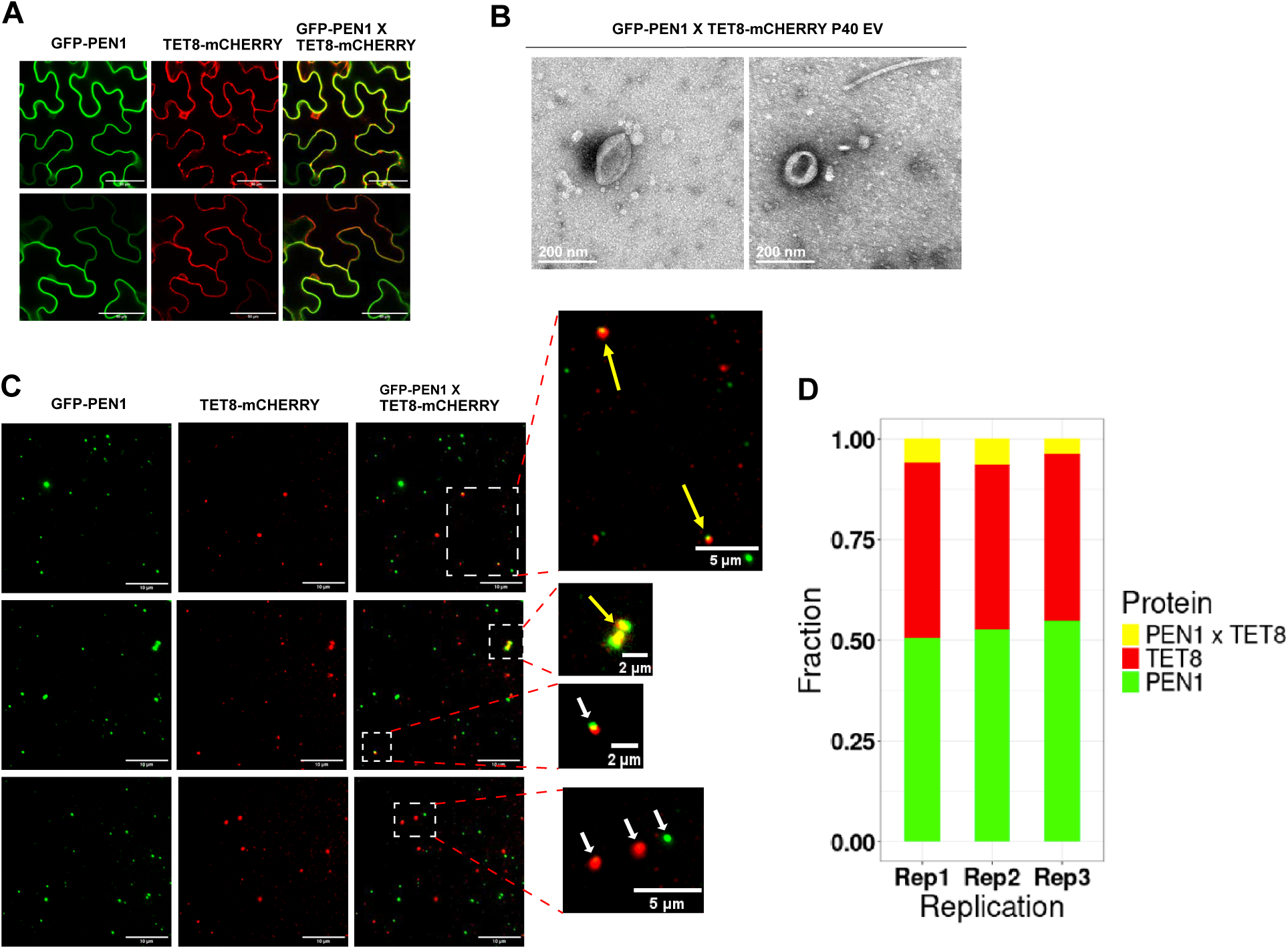
PEN1 and TET8 mark distinct EV populations. **A,** Transient co-expression of PEN1-GFP and TET8-mCherry in pavement cells of *N. benthamiana* leaves. **B,** Negative stain TEM images of P40 EVs isolated from *N. benthamiana* leaves co-expressing these EV markers. **C,** TIRF-M images of P40 EVs isolated from *N. benthamiana* leaves co-expressing PEN1-GFP and TET8-mCherry. The boxed areas are enlarged for better visualization of individual EVs. The Fiji plugin Diana (Schindelin et al., 2012; Gilles et al., 2017) was used to determine colocalization of both proteins. Objects within 250 nm were classified as the same EV (yellow arrows) while objects more than 250 nm apart were classified as different EVs (white arrows). **D**, Stacked bar graph showing co-localization frequency. Scale bars: panel A, 50 µm; panel B, 200 nm; panel C,10 µm; enlarged images of panel C, either 2 µm or 5 µm as marked.

For colocalization of TET8 and PEN1 signals in individual EVs, one potential artifact is aggregation of EVs. To minimize this issue, we used EGTA in our vesicle isolation buffer (VIB) and in our P40 resuspension buffer, which prevents clumping of vesicles (Ebrahimi and Keshtgar, 2020; Borniego et al., 2025). In addition, we used the Fiji plugin Diana (Schindelin et al., 2012; Gilles et al., 2017) to determine colocalization. Objects within 250 nm were classified as the same EV (yellow arrows in Supplementary Figure S4C), while objects more than 250 nm apart were classified as different EVs (white arrows). We chose 250nm as the cut off based on our TEM images and nanoparticle tracking analysis, which show that plant EVs are typically under 250 nm in diameter.

We performed a chi-squared test with the null hypothesis that if proteins sort into vesicles randomly, and the capacity of individual vesicles for proteins is low, vesicles would follow a 1:2:1 ratio of PEN1 alone, PEN1 and TET8, and TET8 alone. A chi-squared test performed on 3 independent replicates (Supplementary Figure S4D) produced a p value less than 0.0001, thus rejecting the null hypothesis, which indicates that PEN1 and TET8 do not sort into vesicles randomly.

Together these results indicate that transient assays in *N. benthamiana* produced results similar to that of transgenic Arabidopsis lines, validating the use of *N. benthamiana* assays for analysis of EV protein sorting. This bypasses the need to create transgenic lines and provides a quick and efficient method to analyze subpopulations of EVs. This also indicates that PEN1 and TET8 are indeed in different vesicle populations. The overlap of 4.7% may be caused, in part, by EV clumping that falls within our 250 nm limit and thus cannot be distinguished from a single EV.

As expected, PEN1 and TET8 rarely co-localized, with an average of just 4.7% of EVs containing both markers across three independent replicates (Figure 2 and Supplementary Figure S4D). This is very similar to that reported by Koch et al. (2024) when analyzing EVs from transgenic Arabidopsis leaves, who found that TET8-GFP and RFP-PEN1 colocalized 7.8% of the time.

**Figure 2.**
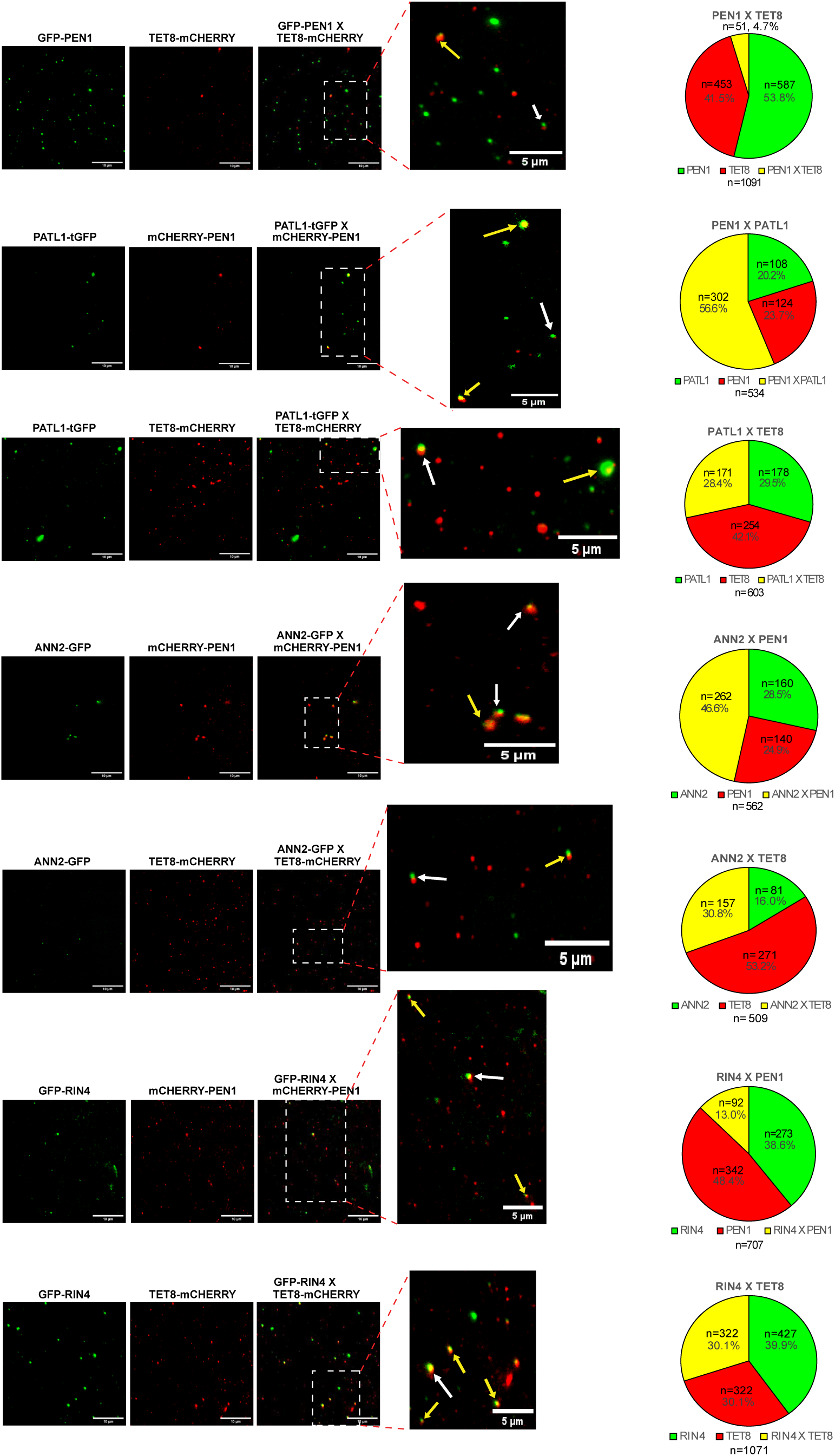
TIRF-M analysis of EV marker co-localization. The indicated EV marker pairs were transiently co-expressed in *N. benthamiana* and then P40 fractions isolated and analyzed by TIRF-M (see Supplementary Figures S4-S10). Puncta were classified by color, with the markers considered to be co-localized when their centers were found to be within 250 nm of each other. Pie charts indicate the fraction of puncta containing individual markers and co-localized markers.

### PATL1 and PEN1 proteins often mark the same EVs

Recent work employing high resolution density gradients has shown that PATL1 and PEN1 co-fractionate, indicating that these proteins may be packaged into the same EV populations (Koch et al., 2024). To test this hypothesis, we transiently co-expressed PATL1 fused with Turbo GFP and PEN1 fused with mCherry in *N. benthamiana* leaves. Supplementary Figure S5A shows the co-expression of both proteins in the epidermal cells of *N. benthamiana*. Most cells expressed both proteins, but not all cells did. We isolated P40 pellets from these leaves and confirmed the presence of EVs using negative stain TEM. Supplementary Figure S5B shows objects with the typical cup-shaped morphology of EVs. PATL1 and PEN1 colocalized in the same EV 56.6% of the time (Figure 2 and Supplementary Figures S5C and S5D). We also performed a chi-squared test with the same null hypothesis as described above, which produced a p value of 0.0063, suggesting that PEN1 and PATL1 also do not sort into vesicles randomly. Since not all cells expressed both proteins, the number 56.6% is likely an underestimate. PATL1 and PEN1 thus appear to be co-packaged into EVs more than would be expected by a random sorting mechanism.

**Supplementary Figure S5.**
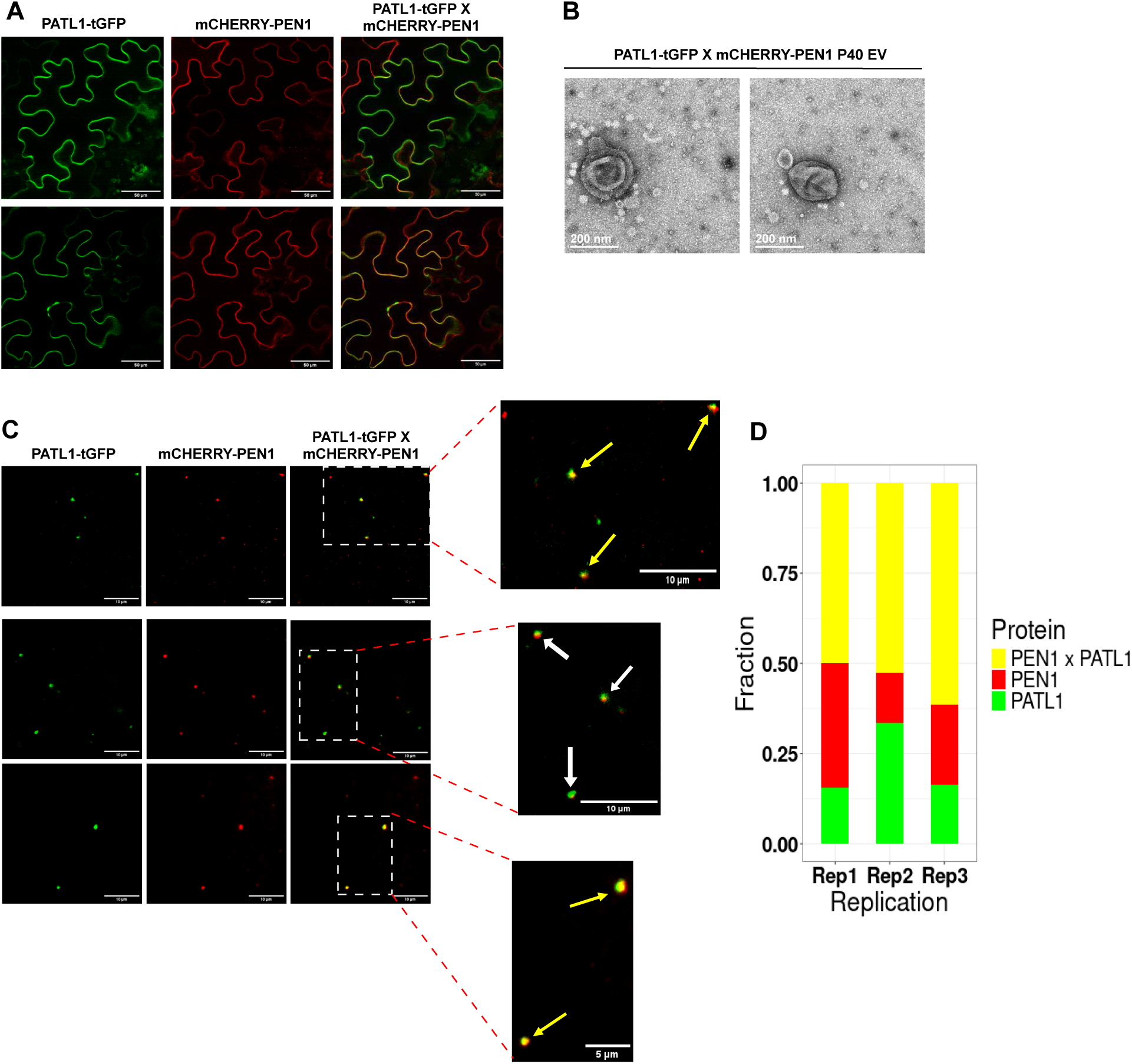
PATL1 and PEN1 frequently co-localize in EVs. **A,** Transient co-expression of PATL1-tGFP and PEN1-mCHERRY in pavement cells of *N. benthamiana* leaves. **B,** Negative stain TEM images of P40 EVs isolated from *N. benthamiana* leaves expressing these EV markers. **C,** TIRF-M images of P40 EVs isolated from *N. benthamiana* leaves co-expressing PATL1-tGFP and PEN1-mCHERRY. The boxed areas are enlarged for better visualization of individual EVs. Co-localization analysis was performed as described in Supplementary Figure S4. **D**, Stacked bar graph showing co-localization frequency. Scale bars: panel A, 50 µm; panel B, 200 nm; panel C,10 µm; enlarged images of panel C, either 5 µm or 10 µm as marked.

### PATL1 and TET8 proteins mark distinct EV populations

Because PATL1 and PEN1 colocalized in EVs the majority of the time, we predicted that PATL1 and TET8 would mark separate EV populations. We co-expressed PATL1-tGFP and TET8-mCHERRY in *N. benthamiana* leaves, and as seen in Supplementary Figure S6A, most cells in *N. benthamiana* expressed both proteins. The TEM images of the P40 fraction confirmed the presence of EVs (Supplementary Figure S6B).

**Supplementary Figure S6.**
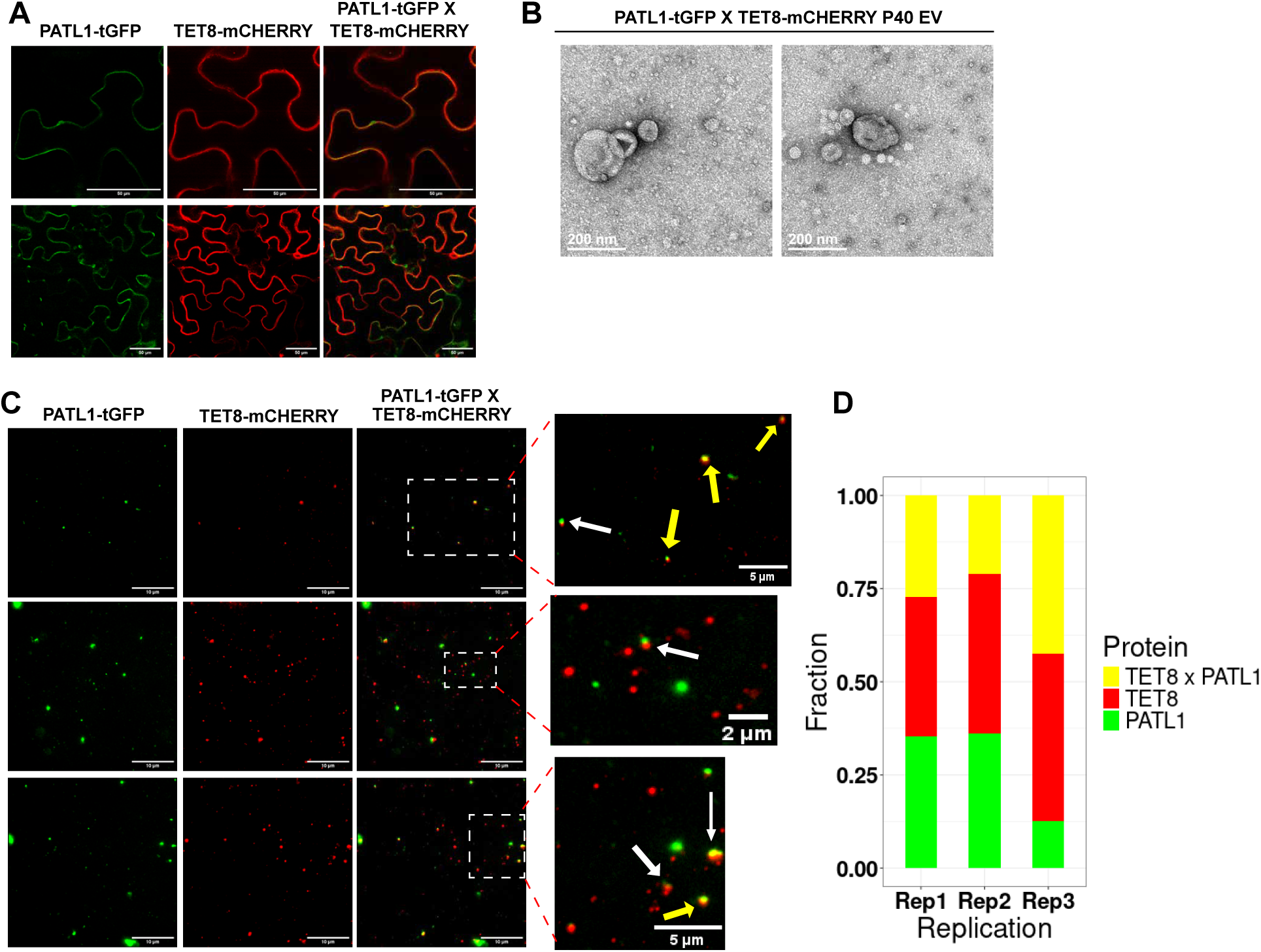
PATL1 and TET8 co-localize in EVs about a third of the time. **A,** Transient co-expression of PATL1-tGFP and TET8-mCHERRY in pavement cells of *N. benthamiana* leaves. **B,** Negative stain TEM images of P40 EVs isolated from *N. benthamiana* leaves co-expressing these EV markers. **C,** TIRF-M images of P40 EVs isolated from *N. benthamiana* leaves co-expressing PATL1-tGFP and TET8-mCHERRY. Co-localization analysis was performed as described in Supplementary Figure S4. **D**, Stacked bar graph showing co-localization frequency. Scale bars: panel A, 50 µm; panel B, 200 nm; panel C,10 µm; enlarged images of panel C, either 2 µm or 5 µm as marked.

We analyzed the P40 fraction using TIRF-M and found that PATL1 and TET8 colocalized 27.2%, 21.1% and 42.5% of the time, (Supplementary Figures S6C and S6D), and the average of the three replicates indicates they colocalized 28.4% of the time (Figure 2). A chi-squared test performed using the null hypothesis described above produced a p value less than 0.0001, indicating that TET8 and PATL1 do not sort randomly into EVs. This result confirms that PATL1 and TET8 are mostly housed in separate EVs; however, they colocalized more often than PEN1 and TET8. Also noteworthy is the fact that TET8 only vesicles are the highest class of vesicles in all three replicates (Supplementary Figure S6D), further supporting our conclusion that TET8-containing EVs are generated by a pathway distinct from PEN1-containing EVs. It also suggests that loading of PATL1 may be somewhat stochastic.

### ANN2 and PEN1 proteins often localize the same EVs

Since PEN1 and TET8 mark different EV populations in both Arabidopsis (He et al., 2021; Koch et al., 2024) and *N. benthamiana*, we wanted to see whether ANN2 and PEN1 localized to separate EVs, which would be expected if ANN2 and TET8 mostly co-localize as previously reported in Arabidopsis (He et al., 2021). We co-expressed ANN2-GFP and mCHERRY-PEN1 in *N. benthamiana* leaves, and as seen in Supplementary Figure S7A, most cells in *N. benthamiana* expressed both proteins. ANN2 appeared to localize to the cytoplasm, while PEN1 mostly localized to the plasma membrane. TEM images confirmed the presence of cup shaped vesicles in the P40 fraction (Supplementary Figure S7B).

**Supplementary Figure S7.**
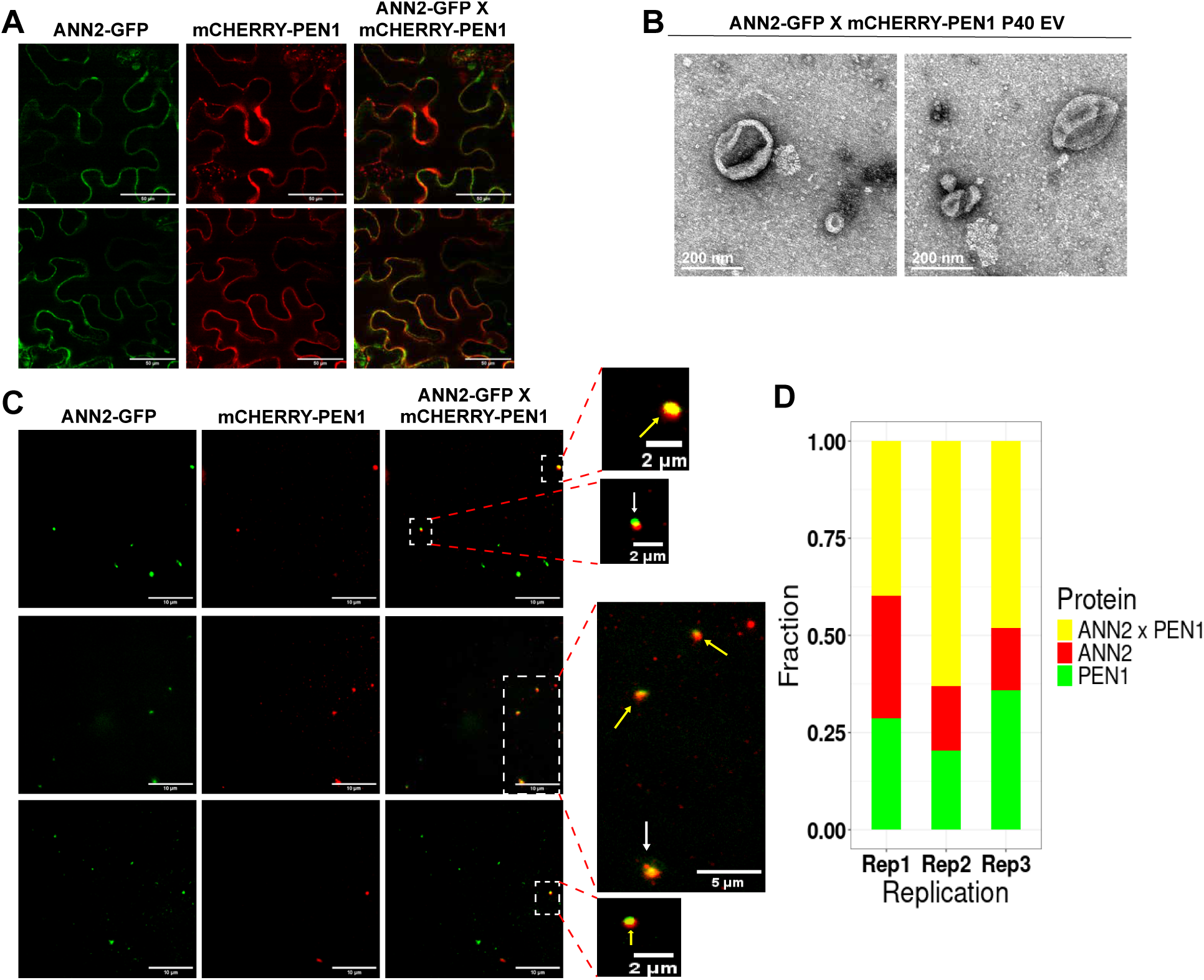
ANN2 and PEN1 are sorted into the same EV about half of the time. **A,** Transient co-expression of ANN2-GFP and PEN1-mCHERRY in pavement cells of *N. benthamiana* leaves **B,** Negative stain TEM images of P40 EVs isolated from *N. benthamiana* leaves co-expressing ANN2-GFP and PEN1-mCHERRY. **C,** TIRF-M of P40 EVs isolated from *N. benthamiana* leaves co-expressing ANN2-GFP and PEN1-mCHERRY. The boxed areas are enlarged for better visualization of individual EVs. Co-localization analysis was performed as described in Supplementary Figure S4. **D**, Stacked bar graph showing co-localization frequency. Scale bars: panel A, 50 µm; panel B, 200 nm; panel C,10 µm; enlarged images of panel C, either 2 µm or 5 µm as marked.

TIRF-M analyses of the P40 fraction revealed that ANN2 and PEN1 co-localized 46.6% of the time (Figure 2 and Supplementary Figure S7D). As with the PEN1 and PATL1 analysis, this may be an underestimate since not all cells expressed both proteins. A chi-squared test produced a non-significant p value of 0.1358, which indicates that the sorting ANN2 and PEN1 is consistent with the null hypothesis that these proteins are sorted randomly into EVs.

### ANN2 and TET8 label distinct EV populations

To confirm that ANN2 and TET8 actually label distinct EVs, we co-expressed ANN2-GFP and TET8-mCHERRY in *N. benthamiana* leaves. As observed with the previous protein pairs, most cells expressed both proteins (Supplementary Figure S8A). TEM imaging confirmed the presence of cup shaped vesicles in the P40 fraction (Supplementary Figure S8B).

**Supplementary Figure S8.**
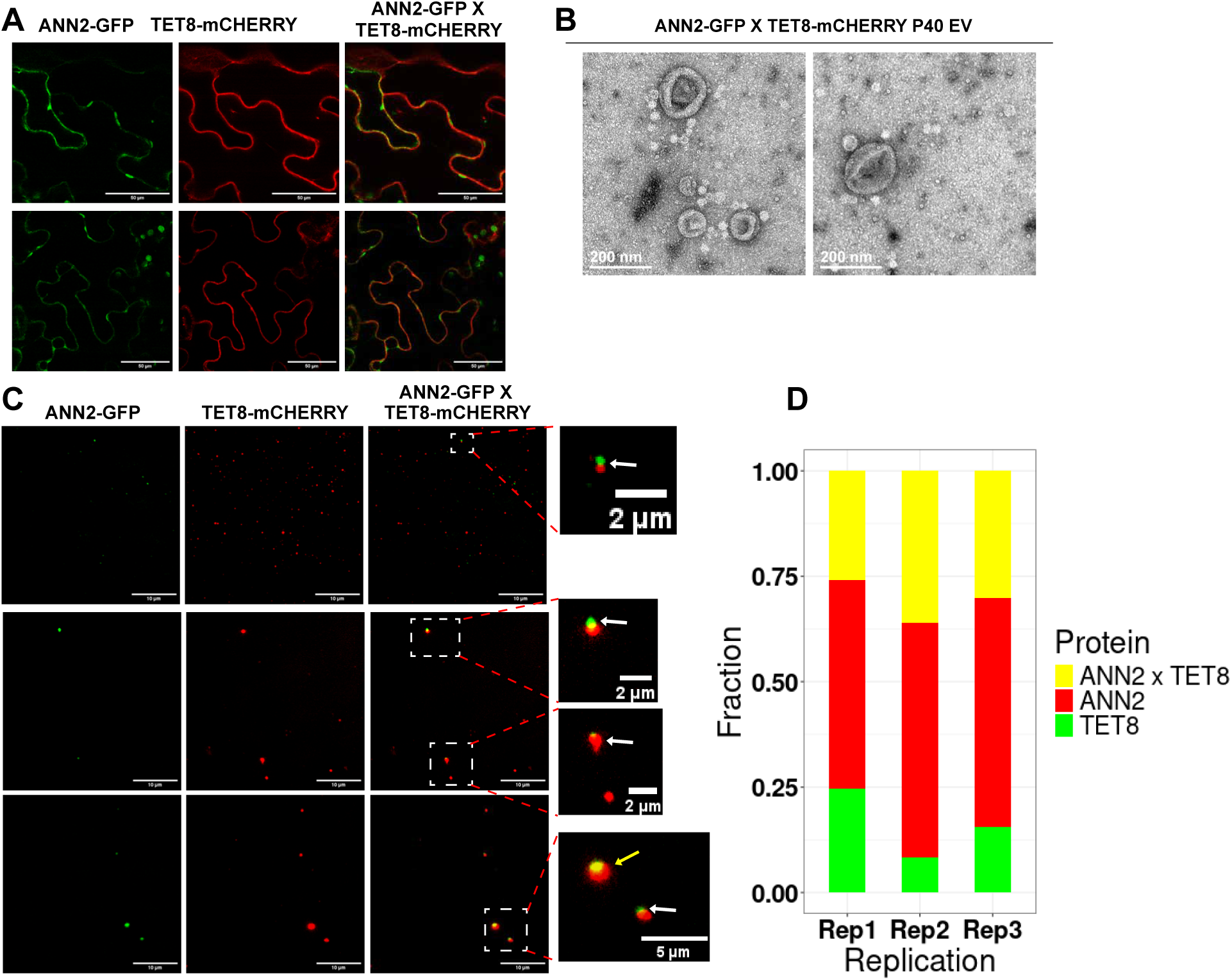
ANN2 and TET8 are sorted in the same EV about one third of the time. **A,** Transient co-expression of ANN2-GFP and TET8-mCHERRY in pavement cells of *N. benthamiana* leaves**. B,** Negative stain TEM images of P40 EVs isolated from *N. benthamiana* leaves co-expressing these EV markers. **C,** TIRF-M of P40 EVs isolated from *N. benthamiana* leaves co-expressing ANN2-GFP and TET8-mCHERRY. Co-localization analysis was performed as described in Supplementary Figure S4. **D**, Stacked bar graph showing co-localization frequency. Scale bars: panel A, 50 µm; panel B, 200 nm; panel C,10 µm; enlarged images of panel C, either 2 µm or 5 µm as marked.

TIRF-M analysis of the P40 fraction revealed that ANN2 and TET8 colocalized 30.8% of the time (Figure 2 and Supplementary Figure S8D). A chi-squared test produced a p value less than 0.0001, indicating that TET8 and ANN2 sorting into EVs is not random. We also observed that TET8-only EVs were the most common class in all three replicates (Supplementary Figure S8D), averaging 53.2% consistent with our conclusion that TET8 marks a special class of EVs. Notably, ANN2-labelled EVs are more likely to be associated with PEN1 than TET8. Together, these results show how diverse plant EVs are in terms of cargo protein content.

### RIN4 co-localizes with TET8 EVs more often than with PEN1 EVs

The last EV marker protein we assessed was RIN4, which is known to associate with EXO70 proteins thus may be involved in regulation of secretion (Redditt et al., 2019). Somewhat unexpectedly, RIN4 co-localized with TET8-containing EVs more frequently than with PEN1-containing EVs when transiently co-expressed in *N. benthamiana* (30% versus 13%; Figure 2 and Supplementary Figures S9 and S10). These findings further support our conclusion that TET8 and PEN1 EVs represent distinct populations likely produced by independent pathways. Notably, however, TET8-only EVs still outnumbered TET8+RIN4 EVs, indicating that RIN4 is not an obligate cargo of TET8 EVs.

**Supplementary Figure S9.**
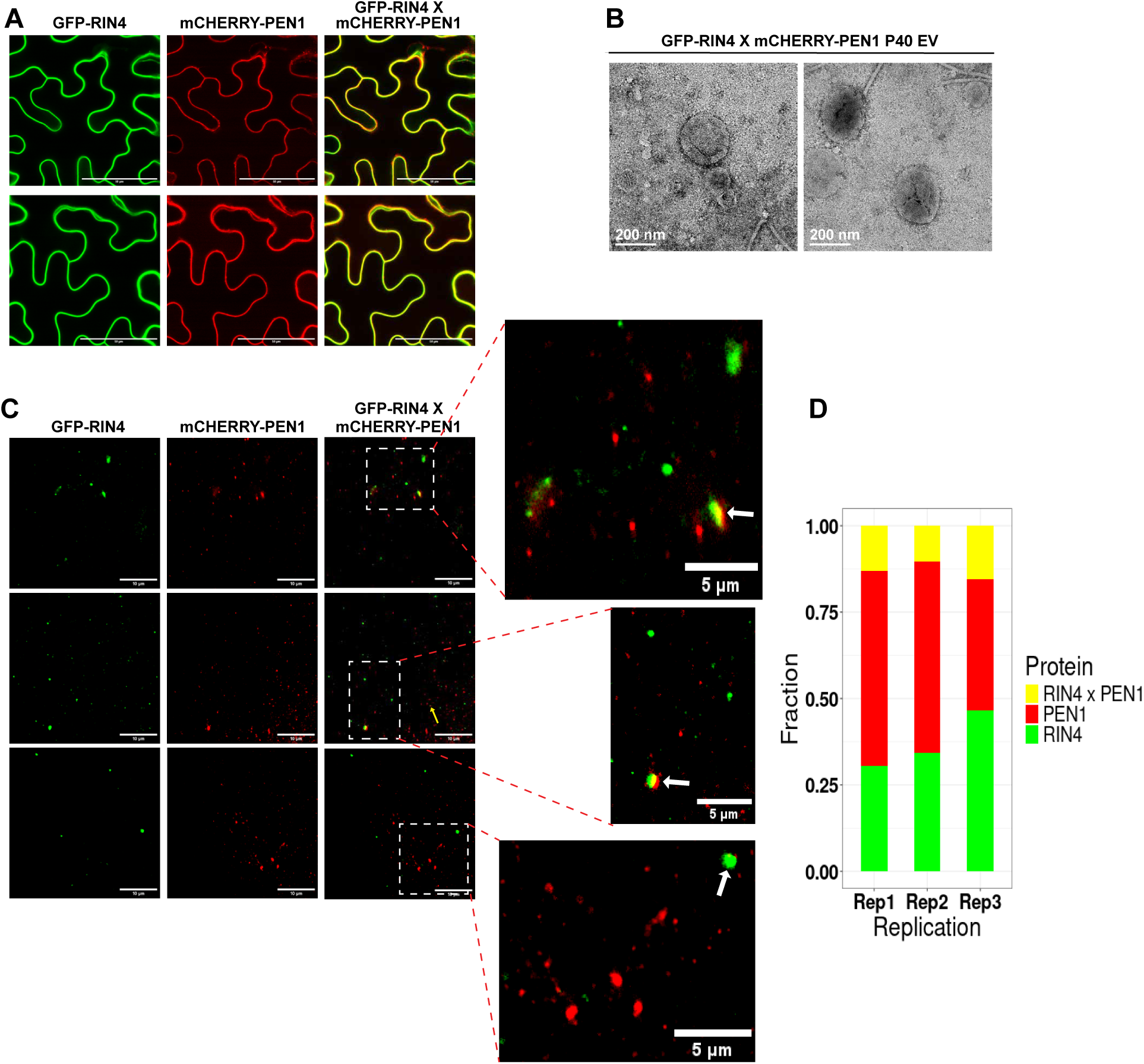
PEN1 and RIN4 rarely sort into the same EV. **A,** Transient co-expression of GFP-RIN4 and mCHERRY-PEN1 in pavement cells of *N. benthamiana* leaves**. B,** Negative stain TEM images of P40 EVs isolated from *N. benthamiana* leaves co-expressing these EV markers. **C,** TIRF-M of P40 EVs isolated from *N. benthamiana* leaves co-expressing GFP-RIN4 and mCHERRY-PEN1. Co-localization analysis was performed as described in Supplementary Figure S4. **D**, Stacked bar graph showing co-localization frequency. Scale bars: panel A, 50 µm; panel B, 200 nm; panel C,10 µm; enlarged images of panel C, 5 µm.

**Supplementary Figure S10.**
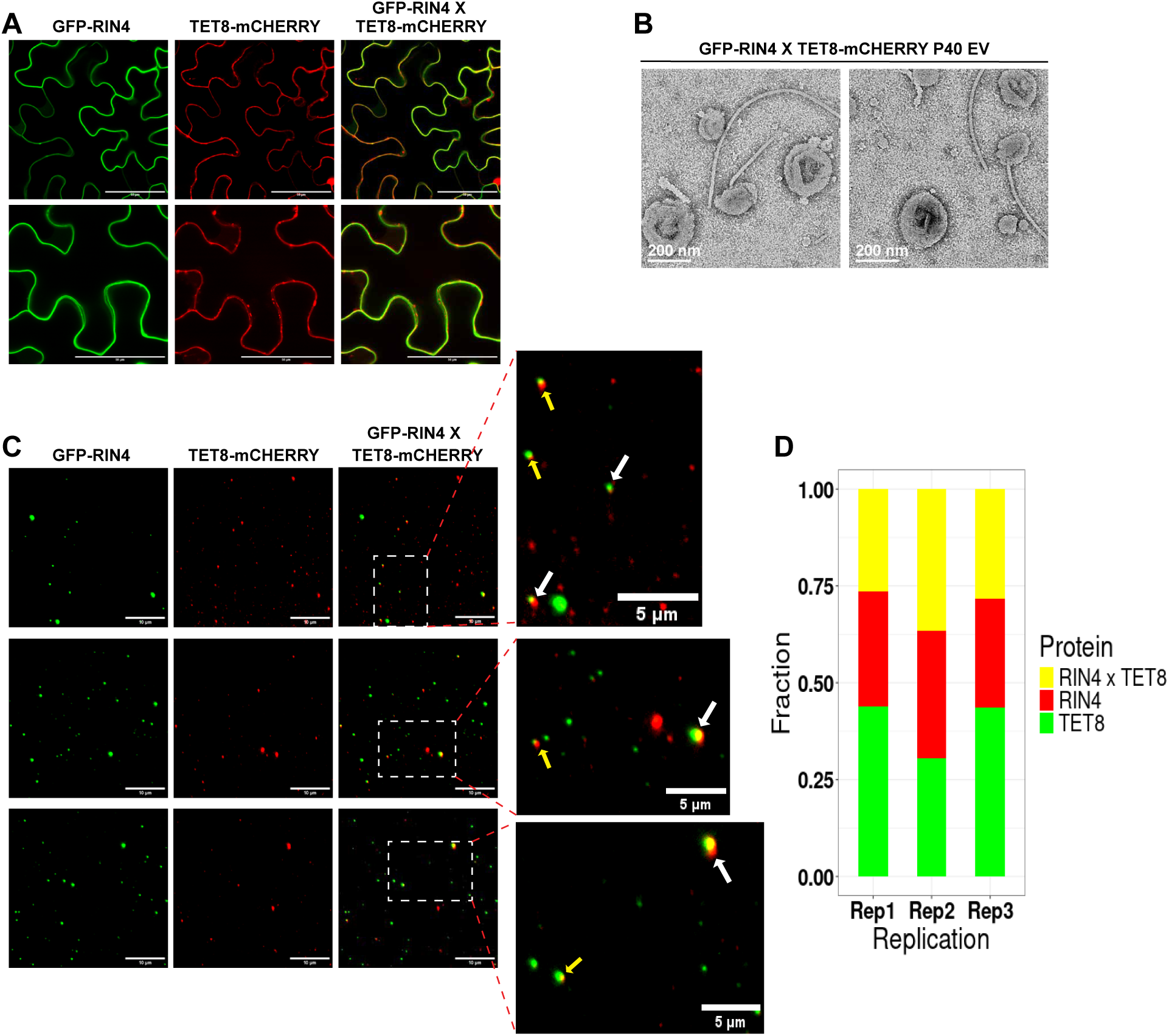
TET8 and RIN4 co-localize in EVs over 30% of the time. **A,** Transient co-expression of GFP-RIN4 and TET8-mCHERRY in pavement cells of *N. benthamiana* leaves**. B,** Negative stain TEM images of P40 EVs isolated from *N. benthamiana* leaves co-expressing these EV markers. **C,** TIRF-M of P40 EVs isolated from *N. benthamiana* leaves co-expressing GFP-RIN4 and TET8-mCHERRY. Co-localization analysis was performed as described in Supplementary Figure S4. **D**, Stacked bar graph showing co-localization frequency. Scale bars: panel A, 50 µm; panel B, 200 nm; panel C,10 µm; enlarged images of panel C, 5 µm.

## Discussion

Studies on mammalian EVs have shown that there are multiple sources of EVs and that EVs are diverse in terms density, protein content, and RNA content (Kowal et al., 2016). Purification of specific subpopulations of EVs can be challenging, however, as density gradient purification is imperfect in separating different subtypes of EVs, which can lead to contradictory results. For example, heat shock protein 70 (HSP70), and major histocompatibility complex I and II proteins were originally thought to mark classic exosomes. However, a recent study by (Kowal et al., 2016) identified them in all EVs. A recent breakthrough came when (Han et al., 2021) used TIRF-M to perform colocalization analysis on multiple markers in individual vesicles, which enabled identification of distinct EV subpopulations.

In this study, we also employed TIRF-M to visualize single vesicles expressing known vesicle markers to identify subpopulations of EVs. We also developed a faster method of studying subpopulations of plant EV markers by using transient expression of EV marker proteins in *N. benthamiana* and quantifying how often they colocalized. This helped us determine which EV markers likely mark the same vesicle populations. We validated this approach by first assessing colocalization of the TET8 and PEN1 markers from Arabidopsis, which have previously been shown to mark distinct vesicle populations (He et al., 2021; Koch et al., 2024). As found in transgenic Arabidopsis, these two proteins are associated with distinct vesicle populations when transiently expressed in *N. benthamiana*.

PEN1 is a SNARE protein that is abundant in the plasma membrane (Fig. 1) (Meyer et al., 2009). Since plant EVs are likely derived, at least indirectly, from plasma membrane, PEN1 may be present in all EVs. Hence, we chose PEN1 as our first choice of EV marker to study. Moreover, PEN1 has previously been shown to be protected inside plant vesicles (Rutter and Innes, 2017; He et al., 2021; Ghosh et al., 2024; Koch et al., 2024). Here, we show that Arabidopsis PEN1 is loaded into EVs when transiently expressed in *N. benthamiana.* Similarly, PATL1 has been shown to be an EV marker protein that is protected inside the lumen of plant vesicles (Rutter and Innes, 2017; Koch et al., 2024). Both PEN1 and PATL1 are upregulated in EVs in response to bacterial infection, suggesting that they play a role in plant immune responses (Rutter and Innes, 2017). PATL1, like PEN1 is present in high, low and medium density Arabidopsis EVs separated using an iodixanol density gradient (Koch et al., 2024). PATL1 binds to phosphoinositides and plays a role in membrane trafficking events (Peterman et al., 2004). A lipid profile of plant EVs found that they are enriched in phosphoinositides as is the plasma membrane, suggesting that PATL1 might be present in all EVs derived from the plasma membrane (Peterman et al., 2004; Suzuki et al., 2016; Liu et al., 2020). We therefore hypothesize that PEN1 and PATL1 may be present together in the majority of plant derived EVs. Indeed, we observed that PEN1 and PATL1 colocalize in the same EV roughly 60% of the time. This finding indicates that PEN1-containing and PATL1-containing EVs likely share the same biogenesis process. Since these proteins were also found in all three densities of EVs, these two proteins might serve as housekeeping proteins that are sorted into EVs by a shared biogenesis method (Meyer et al., 2009; Koch et al., 2024). The observation that EVs can be separated into three different populations based on density argues that there may be three distinct sources of EVs. But the fact that they all are labeled by PEN1 and PATL1 would argue that they share an underlying biogenesis method. There also exists some EVs that only have PEN1, while some that only have PATL1. 60% colocalization might be an underestimate caused by the *N. benthamiana* transient expression system, since not all cells expressed both PEN1 and PATL1. It may also indicate that even though there is an underlying method of EV biogenesis that sorts PEN1 and PATL1 together, whether an individual EV receives both may be stochastic and depends, in part, on the abundance of each protein and the capacity of individual EVs.

TET8 functions as a carrier of sphingolipids, regulating the sorting of glycosyl inositol phosphoceramides (GIPCs) from Golgi bodies to the plasma membrane and eventually to plant EVs (Liu et al., 2024). Deletion of the C terminal tail of TET8 eliminates its ability to bind to GIPCs and its ability to localize to the plasma membrane. Plants expressing this version of TET8 also have reduced levels of GIPCs in the plasma membrane, confirming that TET8 mediates GIPC trafficking (Liu et al., 2024). TET8-containing EVs are thought to carry sRNAs and to be taken up by the necrotrophic fungal pathogen *Botrytis cinera* upon infection of *Arabidopsis*, thus are thought to play a specific role in plant immunity (Cai et al., 2018; He et al., 2021). TET8 also associates with an MVB marker ARA6, which is a Rab5 family GTPase, hence TET8 EVs might also be derived from MVBs (Cai et al., 2018). (He et al., 2021) used transgenic Arabidopsis lines expressing TET8-GFP and mCHERRY-PEN1 to show that TET8 and PEN1 marked different EV populations. Our study confirms that TET8 and PEN1 mark separate EV populations as these proteins colocalized only 5% of the time in P40 pellets. An independent study employing transgenic Arabidopsis used TIRF-M to show that PEN1 and TET8 co-localize in EVs approximately 8% of the time (Koch et al., 2024). Collectively, these studies suggest that there may be a small population of EVs that contain both PEN1 and TET8, or that some clumping of EVs occurs even in the presence of EGTA. On the other hand, PATL1 and TET8 colocalized 31% of the time. This could mean that the PATL1 EVs that do not have PEN1 in them might have TET8 associated with them. This potentially indicates at least three separate EV biogenesis processes, one that sorts PEN1 and PATL1 together in EVs, another that sorts PATL1 and TET8 together in EVs and finally one that sorts only TET8 into EVs.

Annexin2 (ANN2) is an RNA binding protein that binds to sRNA nonspecifically (He et al., 2021). ANN2 is also important for immunity of Arabidopsis against *B. cinerea* (He et al., 2021). ANN2 and TET8 were shown to colocalize in the same puncta when these proteins were transiently expressed in *N. benthamiana* based on the overlap of fluorescence signals (He et al., 2021). However, no measures were taken to minimize EV clumping, which would result in overlapping signal. In addition, this study did not assess whether ANN2 also colocalized with PEN1. It also did not quantify the frequency of vesicles labeled with only ANN2 or only TET8. However, careful analysis of the images shown in this work reveals many vesicles with only ANN2 or TET8. Our results show that ANN2 and PEN1 colocalize more frequently than ANN2 and TET8. The association of ANN2 with PEN1-labeled vesicles suggests that PEN1-labeled vesicles may also carry RNA.

We observed that the frequency of TET8-only EVs was always higher than the frequency of TET8 plus other EV proteins (ANN2, PATL1, and RIN4). In contrast, whenever we expressed PEN1 with a protein other than TET8, the frequency of vesicles expressing both proteins were always higher than EVs expressing PEN1 alone. This is further evidence that TET8-containing vesicles likely have a distinct EV biogenesis pathway. It is notable, though, that RIN4 co-localized more frequently with TET8 than with PEN1 and seemed to be mostly excluded from PEN1-containing EVs. This finding also supports the conclusion that production of PEN1-associated EVs is distinct from production of TET8-associated EVs. Going forward, it will be informative to assess whether RIN4 and/or EXO70 proteins contribute to production of TET8-containing EVs.

Single vesicle imaging of animal EVs has shown that even classical exosome markers such as CD9, CD63, and CD81 do not always colocalize, indicating that the EV population is not only diverse, but that the sorting of EV proteins is somewhat stochastic (Han et al., 2021). This reiterates the importance of analyzing individual EVs using microscopy techniques such as TIRF, along with sample preparation methods that minimize vesicle clumping. TIRF reduces background autofluorescence that can be misinterpreted as vesicle signals as vesicles are small and often in the range of autofluorescence of dust particles on slides. Since the mammalian EV world describes EVs to be heterogeneous, and the sorting of EV proteins to be leaky and stochastic (Han et al., 2021; Fordjour et al., 2022), it is also important to assess the frequency of EV marker co-localization. In both mammalian systems and plant systems, it is not yet clear whether co-localization of specific EV marker proteins affects EV function. Answering this question will require more facile methods for purifying specific EV subpopulations and robust functional assays.

## Materials and Methods

### Plant Growth Conditions

*N. benthamiana* plants were grown in Sun Gro propagation mix (Sun Gro Horticulture, Agawam, MA) under a 16h day and 8h night cycle at 25°C. The soil was supplemented with Osmocote (14-14-14) slow-release fertilizer and watered when dry.

### Plasmid generation for plant expression

PEN1 (<u>AT3G11820</u>) tagged and RIN4 (<u>AT3G25070</u>) tagged with eGFP and TET8 (<u>AT2G23810</u>) tagged with mCherry were cloned into the plant expression vector pEarleyGate100 (pEG100) (www.snapgene.com/plasmids/plant_vectors/pEarleyGate_100) (Earley et al., 2006) using multi-site Gateway cloning (Qi et al., 2012;Reece-Hoyes and Walhout, 2018). This vector places the gene of interest under control of a cauliflower mosaic virus 35S promoter. All three genes were amplified from *Arabidopsis* cDNA. The cDNA was prepared using the Verso cDNA synthesis kit (https://www.thermofisher.com/order/catalog/product/AB1453A) using the manufacturer’s protocol. TET8 was amplified using the primers listed in Table 1, and the resulting PCR product cleaned using a Qiagen PCR clean up kit following the manufactures protocol (https://www.qiagen.com/us/products/discovery-and-translational-research/dna-rna-purification/dna-purification/dna-clean-up/qiaquick-pcr-purification-kit). TET8, was re-amplified using the first PCR product as template using primers with att sites and then cloned into pBSDONR P1-P4 using BP clonase (Qi et al., 2012). PEN1 tagged with eGFP was also cloned into the dexamethasone-inducible expression vector pTA7001-DEST (https://www.addgene.org/71745/) (Gu and Innes, 2011) using multi-site Gateway cloning.

ANN2 (<u>AT5G65020</u>) tagged with eGFP (Qi et al., 2012) was cloned into the plant expression vector pMDC32-HPB (www.addgene.org/32078/) (Qi and Katagiri, 2009) using multi-site gateway cloning. ANN2 was PCR-amplified from an *Arabidopsis* SSP cDNA/ORF Gateway clone (http://signal.salk.edu/SSP/) using primers without att sites. This PCR product was then amplified again using primers with att sites and cloned into pBSDONR-P1P4 (Qi et al., 2012).

PATL1-tGFP was cloned using the Golden Gate cloning method (Werner et al., 2012; Engler et al., 2014). The coding regions of PATL1 (<u>AT1G72150</u>) was amplified from *Arabidopsis* cDNA and the open reading frame was mutated to remove any BsaI restriction sites using site directed mutagenesis, without affecting the open reading frame of the gene. The mutated *PATL1* (<u>AT1G72150</u>) open reading frame was then PCR amplified with appropriate overhangs that matched the promoter and fluorescence tag of the Golden Gate MoClo Plant Parts Kit (www.addgene.org/kits/patron-moclo/#protocols-and-resources) (https://www.addgene.org/kits/marillonnet-moclo/). The promoter Cassava Vein Mosaic Virus (pICSL12006), *PATL1* ORF, Turbo GFP (pICSL50016), and the *Agrobacterium* gene 7 terminator (pICH72400) were assembled using a single digestion event with Bsa1-HF restriction enzyme (New England Biolabs) and T4 DNA ligase, cloning into the Level 1 destination vector (pICH47742) from the MoClo Plant Parts Kit following the protocol provided by New England Biolabs (www.neb.com/en-us/applications/cloning-and-synthetic-biology/dna-assembly-and-cloning/golden-gate-assembly). All cloning fragments were amplified via PCR using Phusion High-Fidelity DNA polymerase (Thermo Scientific). Table 1 shows all sequences of primers used for all cloning steps. The MoClo Plant Parts Kit was a gift from Nicola Patron (Addgene kit # 1000000047) (Engler et al., 2014). The MoClo Toolkit was a gift from Sylvestre Marillonnet (Addgene kit # 1000000044) (Weber et al., 2011; Werner et al., 2012).

### Transient expression in *N. benthamiana*

The above constructs were transformed into *Agrobacterium tumefaciens* strain GV3101(pMP90) and grown on Luria Bertani (LB) agar plates at 28°C supplemented with kanamycin (50 µg/ml) and gentamycin (10 µg/ml) for GFP-PEN1, mCHERRY-PEN1, TET8-GFP and ANN2-GFP, and with kanamycin and carbenicillin (100 µg/ml) for PATL1-tGFP. For transient expression in *N. benthamiana, A. tumefaciens* strains were streaked onto LB agar plates and cells then scraped from these plates 48 hours later and suspended in 10 mM MgCl_2_ and the cell concentration adjusted to an OD_600_=0.6. For co-expression studies, two of the bacterial strains containing the respective expression constructs were mixed 1:1 to obtain a final OD_600_ of 0.3 for each. Acetosyringone (Sigma-Aldrich) was then added to a final concentration of 100 µM and incubated at room temperature for 2.5 h to induce *Vir* gene expression. Bacterial suspensions were then injected into expanding leaves of 4-to-5-week-old *N. benthamiana* plants using a needless syringe. For constructs with a dexamethasone inducible promoter, 25 µM dexamethasone (Thermo Scientific) mixed with 0.02% Tween 20 (Sigma-Aldrich) was sprayed over the leaves to fully wet the leaves at 40 hours post injection.

### Apoplastic wash isolation and EV isolation

4-5-week-old *N. benthamiana* plants were used for all experiments. For constructs expressing EV markers under a constitutive promoter, apoplastic wash fluids were collected 60h post injection of *A. tumefaciens*. For dexamethasone-inducible constructs, apoplastic fluid was collected at 6h, 12h and 24h post dexamethasone application.

To collect apoplastic wash fluid, individual leaves were harvested and vacuum infiltrated with Vesicle Isolation Buffer (VIB; 20 mM MES, 2 mM CaCl_2_, and 0.01 M NaCl, pH 6.0) supplemented with 10 mM EGTA following the protocol described in (Rutter and Innes, 2017). EGTA was included to chelate Ca^2+^ ions present in the VIB and wash fluid, which helps prevent clumping of EVs (Ebrahimi and Keshtgar, 2020; Koch et al., 2024). Vacuum infiltration was performed by submerging the leaves completely in VIB and placing them in a French press coffee maker and applying a vacuum with a vacuum pump (Thermo Savant VP 100 Two Stage Model #1102180403) for approximately 45 seconds until 100% of leaf area was infiltrated. Kimwipes® were used to blot excess buffer from the surface of the leaves. Following that, approximately 5 leaves were placed carefully into a 50 mL needleless syringe, making sure not to damage the leaves. The syringes were then placed in 250 mL wide mouth (33 mm) centrifuge bottles (ThermoFisher catalog number 2103-0008PK). To secure syringes inside the bottles, they were wrapped with parafilm around the top where they met the mouth of the bottles. These assemblies were then centrifuged for 30 mins at 700*g* with a slow acceleration at 4°C using a JA-14 rotor (Avanti J-20 XP centrifuge; Beckman Coulter). Apoplastic wash fluid collected in the bottles was then passed through a 0.22 µm membrane filter to remove debris (e.g., soil particles) and microbes.

For EV isolation, the filtered apoplastic wash fluid was transferred to 13 x 51 mm polycarbonate centrifuge tubes (Beckman Coulter # 349622) and centrifuged at 10,000*g* for 30 mins at 4°C using a TLA100.4 fixed angle rotor and an Optima TLX Ultracentrifuge (Beckman Coulter) to remove any larger cellular contaminants and debris. The resulting supernatants were transferred to fresh 13 x 51 mm polycarbonate centrifuge tubes and centrifuged at 40,000*g* for 1 hour at 4° C. The resulting pellet (the P40) was mixed with fresh VIB supplemented with 10 mM EGTA and centrifuged again at 40,000*g* for 1 hour at 4°C. The final pellet was resuspended in 30 µL VIB supplemented with 10 mM EGTA. For TIRF-M and TEM, EVs were stored at 4°C for a maximum of 2 days prior to microscopy. For protease protection assays and immunoblots, EVs were prepared fresh each time to prevent degradation of EVs upon freeze-thaw.

### Protease protection assay

To assess whether full-length recombinant proteins were located inside the lumens of EVs, a protease protection assay was performed following the protocol described by Rutter and Innes (2017). In brief, P40 EVs were treated with a) Tris-HCl buffer (pH 7.8) as control; b) 1% Triton X100 in Tris-HCl buffer (pH 7.8); or c) 1% Triton X100 for 30 minutes followed by 1 µg/mL trypsin (Promega) for 1.5 hours. All samples were incubated at 37°C for 2 hours before analysis by immunoblotting.

### Protein isolation

For protein isolation from constructs with constitutive promoters, leaf lysate was collected 60h post injection of leaves. Briefly, 500 mg of leaf tissue was frozen using liquid nitrogen and ground using a chilled mortar and pestle followed by the addition of 2 mL of protein extraction buffer (50 mM Tris-HCl pH 7.0, 150 mM NaCl, 0.1% Nonidet P-40, 1% plant protease inhibitor cocktail (Sigma-Aldrich) and 1% 2,2’-dipyridyldisulfide). The resulting suspension was centrifuged at 10,000*g* for 15 mins at 4° C to pellet the debris. The supernatant was collected and used for immunoblots. For dexamethasone inducible GFP-PEN1, leaf lysate was collected from the leaves after apoplastic wash fluid collection at three different time points (6h, 12h and 24h post application of dexamethasone). For dexamethasone inducible free mCHERRY, leaf lysate was collected from the leaves after apoplastic wash fluid collection 12h post application of dexamethasone.

### Immunoblots

For immunoblots of cell lysate and EVs, 12 µL of each were mixed with 3 µL of 5X SDS (sodium dodecyl sulfate) loading buffer (8% w/v SDS in 250 mM Tris-Cl (6.8), 0.1% Bromophenol Blue, 400 mM dithiothreitol and 40% glycerol). The mixture was heated to 95°C for 10 minutes, before loading the samples in 4% to 20% Tris-glycine stain free polyacrylamide gel (BioRad). Gels were run at 180V for 1 hour in Tris/glycine/SDS running buffer. Total proteins were transferred to nitrocellulose membranes (GE Water and Process Technologies) with a constant current of 300 mAmps for 1 hour in TBS buffer (20 mM Tris, 150 mM NaCl, pH 7.5). Protein transfer was confirmed using Ponceau staining. The membrane was blocked with 5% (w/v) Difco Skim Milk (BD Life Sciences) in TBS-T (TBS buffer with 0.1% Tween 20) overnight at 4°C. Membranes were washed with TBS-T buffer again before incubating with primary antibodies for 2 hours. The primary antibodies were all mixed with 5% (w/v) milk in TBS-T buffer using the following dilutions: anti-GFP (Sigma-Aldrich 66002-1-Ig) (1:2000); anti-RFP (Chromotek 6G6) (1:2000); anti-PATL1 (Peterman et al., 2004) (1:1000). After primary antibody treatment, membranes were washed with TBS-T five times for 10 mins each and then incubated for 2 hours in secondary antibody mixed in 5% milk in TBS-T buffer using the following dilutions: anti-mouse-HRP (Abcam AB6789) (for anti-GFP and anti-RFP) 1:5000; anti-rabbit-HRP (Abcam AB97051) (for anti-PATL1) 1:5000. Blots were then washed in TBST-T five times for 15 mins each and then the membranes were imaged using Super Signal West Femto Maximum Sensitivity Substrate (Thermo Scientific) and a BIO-RAD ChemiDoc imaging system.

### Transmission Electron Microscopy

To image EVs by transmission electron microscopy (TEM), approximately 5 µL of P40 suspension was spotted onto a glow discharged (15 mA for 60 seconds) Formvar and carbon coated Cu electron microscopy grid (Electron Microscopy Sciences). After 10 mins, the excess sample was wicked off using a clean filter paper. Following that,10 µL of 2% (w/v) uranyl acetate was placed on the grid and allowed to incubate for 5 mins. The excess uranyl acetate was wicked off using a filter paper and the grid was allowed to dry overnight. The grids were then imaged using a JOEL JEM 1010 transmission electron microscope (JOEL USA) at 80 kV.

### Confocal microscopy

To determine the expression and subcellular localization of fluorescently tagged EV marker fusion proteins, *N. benthamiana* leaves were imaged using a Leica Stellaris 8 confocal microscope fitted with a 63x water immersion objective. For non-inducible constructs (PATL1-tGFP, mCHERRY-PEN1, TET8-mCHERRY, GFP-PEN1 and ANN2-GFP, pICH47742 empty vector, pEG100 empty vector and pMDC32 empty vector) leaves were imaged 60h after infiltration. For the dexamethasone inducible GFP-PEN1 construct, and its empty vector control (pTA7001), leaves were imaged at 3 time points post dexamethasone spray (6h, 12h and 24h). Each construct was imaged at least three times from separate infiltration events. Leaves infiltrated with 10 mM MgCl_2_ alone were also imaged as negative controls for each imaging experiment. To image GFP and tGFP fusions, we used an excitation wavelength of 488 nm and an emission wavelength between 500nm and 540 nm. To image mCHERRY fusions, we used an excitation wavelength of 587 nm and an emission wavelength between 600 and 640 nm. Differential Interference Contrast (DIC) images were taken in parallel to fluorescence images. Images were auto corrected for brightness and contrast using Fiji (Schindelin et al., 2012).

### Total Internal Reflection Fluorescence Microscopy and Fiji data analysis

To avoid freeze-thaw events, for all TIRF-M imaging, we used fresh P40 EV pellets that were either isolated the same day or kept at 4°C for no longer than 48 h. These P40 pellets were resuspended in VIB supplemented with 10 mM EGTA to prevent clumping of EVs. Slides (22 x 22 mm) and cover slips (1.5) were cleaned with 95% ethanol several times using a Kimwipe and then allowed to dry overnight. For TIRF-M, 8 to 10 µL of P40 EV suspension was placed in the middle of the slide and the coverslip was placed carefully and evenly to avoid bubbles. The cover slip was sealed in place with clear nail-polish and imaged using a GE DeltaVision OMX Super Resolution (SR) microscope and a TIRF lens using TIRF settings and 1.516 refractive index immersion oil. The focal plane was found at roughly 6700 ± 100 µm by adjusting the z-direction in 5 µm intervals and then switching to 0.5 µm intervals to fine tune the focus on EVs. The laser setting was kept the same for control and fluorescent EVs, and care was taken to change the xy position of the slide for each picture to avoid photobleaching of EVs.

The Fiji plugin DiAna (Distance Analysis) was used to measure the distance between two colocalized objects (Gilles et al., 2017) (https://imagej.net/plugins/distance-analysis). The radial center of the fluorescent objects was determined using gaussian methods. Any particles whose radial centers were more than 250 nm apart were considered not colocalized. This distance was used because most plant EVs are less than 250 nM in diameter (Rutter and Innes, 2017), thus two objects more than this distance apart are unlikely to be contained in a single EV. Multiple images were analyzed for each combination of EV markers. The percentage colocalization was calculated using a Microsoft Excel spreadsheet.

## ACKNOWLEDGEMENTS

We thank Barry Stein and the IU Bloomington Electron Microscopy Center at Indiana University for access to a JEOL JEM 1010 Transmission Electron Microscope. We are also grateful for the assistance of Benjamin Koch and Andras Kun with TIRF microscopy and the IU Light Microscopy Imaging Center for access to a Leica SP8 confocal microscope GE Deltavision OMX SR super-resolution microscope. We thank Benjamin Koch, Jessica Foster and Thomas Redditt for helping with the cloning. We also thank Brian Rutter for developing the vesicle isolation protocol in the Innes lab. Finally, we would like to thank Paul Christopher Hunt for his help with the statistical analysis. The IU Light Microscopy center’s purchase of the GE Deltavision OMX SR super-resolution microscope was supported by a grant from the National Institute of Health (grant number NIH1S10OD024988-01). This project was supported by grants from the United States National Science Foundation Plant Biotic Interactions and Plant Genome Research programs (grant numbers IOS-1645745, IOS-1842685, and IOS-2141969 to R.W.I.). S.G. was supported by a Carlos Miller fellowship from the Indiana University Foundation.

---

**Supplementary Table S1.**
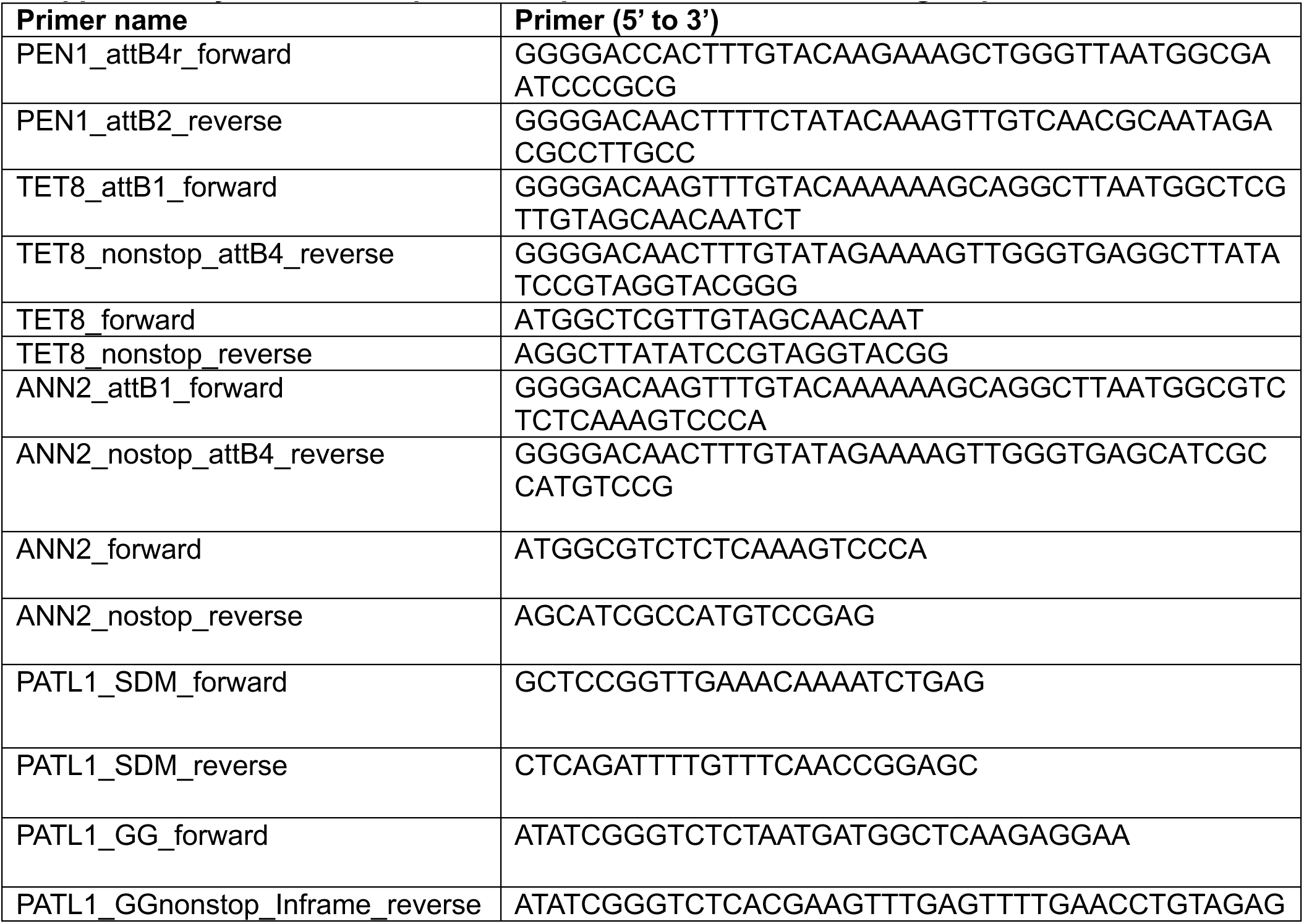
Sequences of primers used for all cloning steps.

